# Chronic inflammation suppresses del(5q)-like MDS HSCs via p53

**DOI:** 10.1101/2022.06.22.497183

**Authors:** Tomoya Muto, Callum S. Walker, Kwangmin Choi, Madeline Niederkorn, Chiharu Ishikawa, Melinda Varney, Kathleen Hueneman, Daniel T. Starczynowski

**Author notes:** **Corresponding author:** Daniel Starczynowski, Division of Experimental Hematology and Cancer Biology, Cincinnati Children’s Hospital Medical Center, Cincinnati, OH, USA.

## Abstract

Inflammation is associated with the pathogenesis of Myelodysplastic syndromes (MDS). Emerging evidence suggests that MDS hematopoietic stem and progenitor cells (HSPCs) exhibit an altered response to systemic low-grade inflammation, which contributes to their competitive advantage over wild-type HSPCs and ensuing hematopoietic defects. Deletion of the long arm of chromosome 5 (del(5q)) is the most common chromosomal abnormality in patients with MDS. Although this subtype of MDS contains several haploinsufficient genes that directly impact innate immune signaling, the effects of an inflammatory milieu on del(5q) MDS HSPCs remains poorly defined. Utilizing a model of del(5q)-like MDS, wherein two 5q genes, miR-146a and TIFAB, are deleted, we found that chronic low-grade inflammation impaired the function of del(5q)-like MDS HSPCs and contributed to a more severe disease. The del(5q)-like MDS HSPCs exposed to chronic inflammation became less quiescent, but without changes in cell viability. In response to inflammation, mouse and human del(5q) MDS HSPCs activated a partial p53 response. The impaired function and reduced cellular quiescence of del(5q) MDS HSPCs exposed to inflammation could be restored by deletion of p53. Since TP53 mutations are highly enriched in del(5q) AML patients following an initial MDS diagnosis, increased p53 activation in del(5q) MDS HSPCs due to inflammation may create a selective pressure for genetic inactivation of p53. These findings uncover the contribution of systemic inflammation on dyshematopoiesis in del(5q) MDS and provide a potential explanation for acquired p53 mutations in myeloid malignancies with del(5q).

## Introduction

Myelodysplastic syndromes (MDS) are clonal hematopoietic stem and progenitor cell (HSPC) malignancies defined by blood cytopenia, myeloid cell dysplasia, and ineffective hematopoiesis^1,2^. Hemizygous deletion of the long arm of chromosome 5q (del(5q)) is the most common cytogenetic alteration in MDS. Del(5q) is also found in 25% of therapy-related MDS cases and is strongly correlated with progression to AML^3-5^. Patients with a sole deletion of 5q exhibit macrocytic anemia, leukopenia, and increased platelets. Bone marrows (BM) from these patients show dysplastic mononuclear megakaryocytes, erythroid hypoplasia, and low blasts (<5%). Another underlying feature of MDS, including del(5q) MDS, is the contribution of systemic inflammation. The MDS BM niche exhibits increased inflammatory signaling, including elevated cytokines, chemokines, and alarmins. Microenvironmental alterations are partly due to aging, but there is also evidence that MDS hematopoietic cells themselves alter the stem-cell niche through activation of the innate immune system and related inflammatory signaling. Recently, several studies have implicated that low-grade systemic inflammation favors the expansion of pre-leukemic or MDS HSCs over normal HSCs^6-10^. Thus, systemic inflammation in MDS has pleotropic effects on the etiology of the disease.

Contiguous deletions of chromosome (chr) 5q span several megabases encompassing many critical genes. Over the past several years, haploinsufficiency of several chr 5q genes have been implicated in del(5q) MDS, including genes that play important roles in ribosome function (RPS14), cell cycle regulation (CDC25c/PP2A), innate immune signaling (miR-145, miR-146a, TIFAB, and DIAPH), stress response signaling (EGR1), and β-catenin signaling (APC, CSNK1A1)^11^. Deletion of RPS14 results in a p53-dependent defect in erythropoiesis, but not in a competitive advantage of HSPCs^12^. Moreover, loss of RPS14 coincides with increased expression of alarmins and non-cell autonomous altered innate immune signaling^13,14^. Deletions of CSNK1A1, EGR1, and APC are implicated in the proliferation of hematopoietic cells. Among these genes, genetic barcoding revealed that CSNK1A1 haploinsufficiency is the major driver of clonal expansion of del(5q) MDS HSPCs^15^. CSNK1A1 and APC are both regulators of β-catenin function, which is thought to contribute to stem cell-self renewal programs in del(5q) HSCPs. Deletion of miR-145, miR-146a, mDia1, and TIFAB collectively contribute to increased innate immune signaling in MDS HSPCs, albeit through distinct mechanisms^16-18^. Deletion of mDia1, a RhoA GTPase effector that belongs to the formin protein family, increases innate immune signaling in neutrophils due to upregulation of CD14 and results in an MDS-like phenotype in mice^19^. Deletion of miR-145, results in derepression of Mal/TIRAP, which is important for an initial step of signaling by the Toll-like receptor (TLR) superfamily^20,21^. Deletion of miR-146a increases TRAF6 and IRAK1 mRNA and translation, while loss of TIFAB increases TRAF6 protein stability, thus resulting in overexpression and activation of TRAF6 and IRAK1 in MDS HSCs in the absence of ligand-mediated activation of the TLRs^21-29^. TIFAB is a fork-head associated domain protein that induces lysosomal degradation of TRAF6 protein, while deletion of TIFAB results in increased TRAF6 protein expression^25,30^. Reduced TIFAB expression also results in diminished USP15 de-ubiquitinase function and consequently p53 activation in hematopoietic cells^31^. Deletion of TIFAB in mouse hematopoietic cells results in modest hematopoietic defects, and occasionally BM failure^25^. However, concurrent hematopoietic-specific deletion of TIFAB and miR-146a results in higher levels of TRAF6 expression and innate immune pathway activation, which coincides with a highly penetrant BM failure, a diseased phenotype that more faithfully recapitulates human del(5q) MDS^24^.

Despite overwhelming evidence of chronic inflammation in the BM and PB of del(5q) MDS patients, the effects of systemic inflammatory signals on del(5q) MDS HSPCs and disease progression remains uncharacterized and under ongoing investigations. In this study, we focused on the effects of systemic low-grade inflammation in del(5q) MDS by utilizing a mouse model in which miR-146a and TIFAB, two critical mediators of aberrant innate immune signaling in MDS HSPCs, are co-deleted. We found that low-grade inflammation accelerates the del(5q) MDS-like phenotype and provide a potential explanation for the high rate of TP53 mutations in patients with del(5q).

## Results

### Low-grade inflammation induces a bone marrow failure in *Tifab*^−/-^;*miR-146a*^−/-^ mice

To investigate the impact of a chronic low-grade inflammation on the pathogenesis of del(5q) MDS, we utilized a mouse model in which Tifab and miR-146a were simultaneously deleted (*Tifab*^*-/-*^*;miR-146a*^*-/-*^), as previously described^24^. TIFAB and miR-146a are co-deleted in ∼80% of del(5q) MDS patients and are both implicated in innate immune and inflammatory signaling. An inflammatory milieu was established using the gram-negative bacterial component lipopolysaccharide (LPS), a ligand for TLR4, which induces systemic inflammation and affects HSPC function^32^. Chronic low-dose (LD) treatment with LPS (1 μg/g, hereafter LD-LPS) was administered via intraperitoneal (i.p.) injection twice a week for 30 days into *Tifab*^−/-^;*miR-146a*^−/-^ or wild-type (WT) mice (**Fig. 1A**). After the last dose of LD-LPS or vehicle, hematopoietic cell chimeric mice were then established by transplanting bone marrow (BM) cells from the treated *Tifab*^−/-^;*miR-146a*^−/-^ or WT mice into lethally-irradiated CD45.1 WT mice. As we reported previously^24^, mice engrafted with *Tifab*^−/-^;*miR-146a*^−/-^ BM cells developed multilineage cytopenias, including neutrophilia, anemia and thrombocytopenia 4 months post-transplant as compared to mice engrafted with WT BM cells (**Fig. 1B**). As expected, the severity of cytopenias following LD-LPS administration in mice engrafted with *Tifab*^−/-^;*miR-146a*^−/-^ BM cells was worse as compared to LD-LPS treated mice engrafted with WT BM cells (**Fig. 1B**). Surprisingly, LD-LPS administration did not contribute to more severe cytopenias in mice engrafted with *Tifab*^−/-^;*miR-146a*^−/-^ BM cells as compared to untreated mice (**Fig. 1B**). Mice engrafted with WT BM cells exhibited significant differences in PB counts (neutrophils, lymphocytes, and platelets) as compared to mice treated with LD-LPS (**Fig. 1B**). The proportion of myeloid cells (CD11b+Gr1-) and lymphoid cells (CD3+ and B220+) in the PB were significantly reduced following LD-LPS treatment in mice engrafted with *Tifab*^−/-^;*miR-146a*^−/-^ BM cells as compared to WT mice (**Fig. 1C**).

**Fig. 1.**
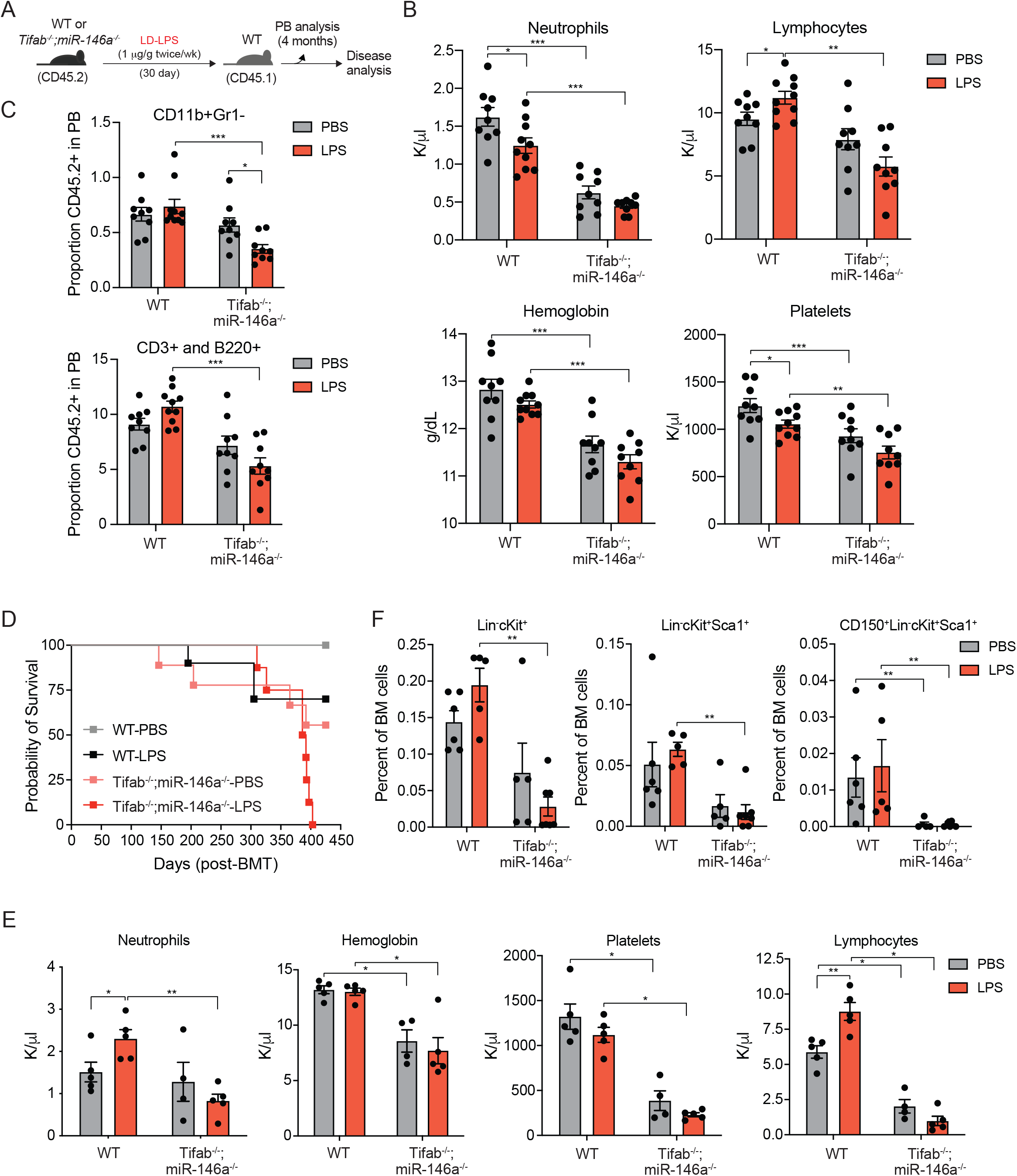
LD-LPS induces cytopenias and accelerated disease in Tifab^−/-^;miR-146a^−/-^ mice. **(A)** Outline of BM transplantations using WT or Tifab^−/-^;miR-146a^−/-^ mice in the presence of low-dose LPS (LD-LPS). **(B)** PB counts of the recipient mice at 4 months after transplantation (n = 9-10 per group). **(C)** Summary of lymphoid (CD3+ and B220+), myeloid (CD11b+Gr1-) proportions within the PB of the recipient mice (n = 9-10 per group). **(D)** Overall survival of mice transplanted with WT or Tifab^−/-^;miR-146a^−/-^ mice treated with either PBS or LPS (n = 9-10). **(E)** PB counts of the moribund Tifab^−/-^;miR-146a^−/-^ mice (n = 4-5 per group) at 10-13 months after transplantation. **(F)** Proportion of the indicated hematopoietic myeloid progenitors in the BM of 2 months after transplantation (n = 5-7 per group). Significance for panels B, C, E, and F was determined with a Student’s *t* test (*, *P* < 0.05; **, *P* < 0.01; ***, *P* < 0.001).

Deletion of Tifab and miR-146a results in activation of IRAK1 and TRAF6 signaling^24^. Therefore, to determine whether increased TRAF6-IRAK signaling contributes to cytopenias in *Tifab*^−/-^;*miR-146a*^−/-^ mice, we utilized an IRAK1/4 inhibitor (NCGC-1481) in vivo^33^. As above, BM cells from *Tifab*^−/-^;*miR-146a*^−/-^ or WT mice were transplanted into lethally-irradiated CD45.1 WT mice. Once the *Tifab*^−/-^;*miR-146a*^−/-^ recipient mice developed PB cytopenias (∼20 weeks post engraftment), the WT and *Tifab*^−/-^;*miR-146a*^−/-^ recipient mice were randomized and treated daily for 2 weeks with 30 mg/kg of the IRAK1/4 inhibitor or vehicle (PBS), as previously described^33,34^ (**Supplemental Fig. 1A**). The *Tifab*^−/-^;*miR-146a*^−/-^ recipient mice treated with PBS progressively developed anemia and neutropenia (**Supplemental Fig. 1B**). However, treatment of the *Tifab*^−/-^;*miR-146a*^−/-^ recipient mice with the IRAK1/4 inhibitor resulted in improved PB counts (**Supplemental Fig. 1B**), suggesting that the cytopenias associated with deletion of TIFAB and miR-146a in del(5q) MDS are partly attributed to increased IRAK-TRAF6 signaling.

To evaluate the effect of a chronic low-grade inflammation on disease development, we monitored the mice for over 1 year and found that all mice engrafted with *Tifab*^−/-^;*miR-146a*^−/-^ BM cells that were treated with LD-LPS (10/10) succumbed to disease by 400 days (**Fig. 1D**). In contrast, approximately 50% (5/11) of mice engrafted with *Tifab*^−/-^;*miR-146a*^−/-^ BM cells treated with vehicle (PBS) survived up to 425 days. The majority of WT mice (7/10) treated with LD-LPS were also alive at the conclusion of the experiment (**Fig. 1D**). The moribund mice transplanted with *Tifab*^−/-^;*miR-146a*^−/-^ BM cells that were treated with PBS or LD-LPS developed multi-lineage cytopenias, reduced BM cellularity, and splenomegaly, which are consistent with findings of BM failure. (**Fig. 1E, Supplementary Fig. 2A-D**). However, administration of LD-LPS resulted in a more severe cytopenias and evidence of BM failure at time of death (**Fig. 1E**). Examination of the hematopoietic stem and progenitor cells showed that *Tifab*^−/-^;*miR-146a*^−/-^ BM cells produce fewer HSCs (CD150+Lin-cKit+Sca1+) and HPCs (Lin-cKit+Sca1+ and Lin-cKit+) than WT BM cells. Treatment with LD-LPS resulted in a further reduction in *Tifab*^−/-^;*miR-146a*^−/-^ HSCs and HPCs (**Fig 1F**). In contrast, LD-LPS induced an expansion of WT HPCs as expected based on previous studies. Altogether, these observations indicate that chronic low-grade inflammation does not meaningfully contribute to worse cytopenias but can accelerate disease development in a del(5q)-like MDS model.

### *Tifab*^*-/-*^*;miR-146a*^*-/-*^ BM HSPCs exhibit a competitive advantage following low-grade inflammation

MDS HSPCs co-exist with WT cells in the BM, and recent studies have demonstrated MDS HSPCs outcompete WT HSPCs primarily under conditions of chronic inflammation^6-10^. This suggest that low-grade inflammation in MDS patients is a key determinant of the relative competitive advantage of MDS HSPCs. Therefore, to examine the relative competitive advantage of del(5q)-like HSPCs to WT HSPCs in an inflammatory milieu, we performed *in vivo* BM cell competition assays using equal mixture of WT and *Tifab*^*-/-*^*;miR-146a*^*-/-*^ BM cells. To avoid a differential response of CD45.2- and CD45.1-expressing hematopoietic cells to LPS as reported^35^, we utilized CD45.2 WT mice expressing GFP in under the control of the UBC promoter (WT-GFP^36^), which could be distinguished from CD45.2 *Tifab*^*-/-*^*;miR-146a*^*-/-*^ BM cells (GFP-). We co-transplanted BM cells from *Tifab*^*-/-*^*;miR-146a*^*-/-*^ mice and UBC-GFP transgenic mice (one to one ratio) into lethally irradiated CD45.1 WT mice. Two months post-engraftment, chimeric mice were injected i.p. with LD-LPS or PBS twice a week for 4 weeks (**Fig. 2A**). At this timepoint (month 3 post transplant), we observed a trending decrease in the relative frequency of CD45.2^+^GFP^−^ *Tifab*^*-/-*^*;miR-146a*^*-/-*^ cells in the blood of chimeric mice treated with LD-LPS compared to PBS-treated chimeric mice (**Fig. 2B**). Examination of HSPCs in the BM at month 3 also revealed reduced proportions of CD45.2^+^GFP^−^ *Tifab*^*-/-*^*;miR-146a*^*-/-*^ LK and LSK cells compared to that of CD45.2^+^GFP^+^ WT cells of chimeric mice treated with LD-LPS compared to PBS-treated chimeric mice (**Fig. 2C**,**D**). Next, BM cells from the chimeric mice were isolated and serially transplanted into lethally-irradiated CD45.1 WT mice, followed by evaluation of the proportions of GFP^+^ WT cells and GFP^−^ *Tifab*^*-/-*^*;miR-146a*^*-/-*^ cells among total PB cells every month for 3 months post each of the transplantations (**Fig. 2B**). The proportion of CD45.2^+^GFP^−^ *Tifab*^*-/-*^*;miR-146a*^*-/-*^ cells derived from the PBS-treated mice decreased from 70% at month 3 to 35% at month 4 after the secondary BM transplantation (**Fig. 2B**). Although proportions of CD45.2^+^GFP^−^ *Tifab*^*-/-*^*;miR-146a*^*-/-*^ cells from LD-LPS-treated mice also declined after the secondary BM transplantation, the proportion of the *Tifab*^*-/-*^*;miR-146a*^*-/-*^ cells were consistently higher as compared to the PBS-treated mice (**Fig. 2B**). Interestingly, the proportion of CD45.2^+^GFP^−^ *Tifab*^*-/-*^*;miR-146a*^*-/-*^ cells among LKs, LSKs and CD150^+^CD48^−^LSKs (HSCs) in the BM significantly decreased from ∼60% (month 3) to ∼20% (month 9) relative to the GFP^+^ WT cells upon secondary BM transplantation of cells derived from PBS-treated mice (**Fig. 2C**,**D**), indicating that del(5q)-like MDS HSCs are functionally less competitive than WT HSCs without exposure to inflammation. In contrast, the proportion of CD45.2^+^GFP^−^ *Tifab*^*-/-*^*;miR-146a*^*-/-*^ LKs, LSKs and HSCs derived from the LD-LPS mice increased from ∼50% (month 3) to ∼75% (month 7) upon secondary BM transplantation (**Fig. 2C**,**D**). These findings suggest that del(5q)-like MDS HSCs are functionally impaired but sustain a long-term competitive advantage compared to WT HSCs cells following exposure to low-grade chronic inflammation.

**Fig. 2.**
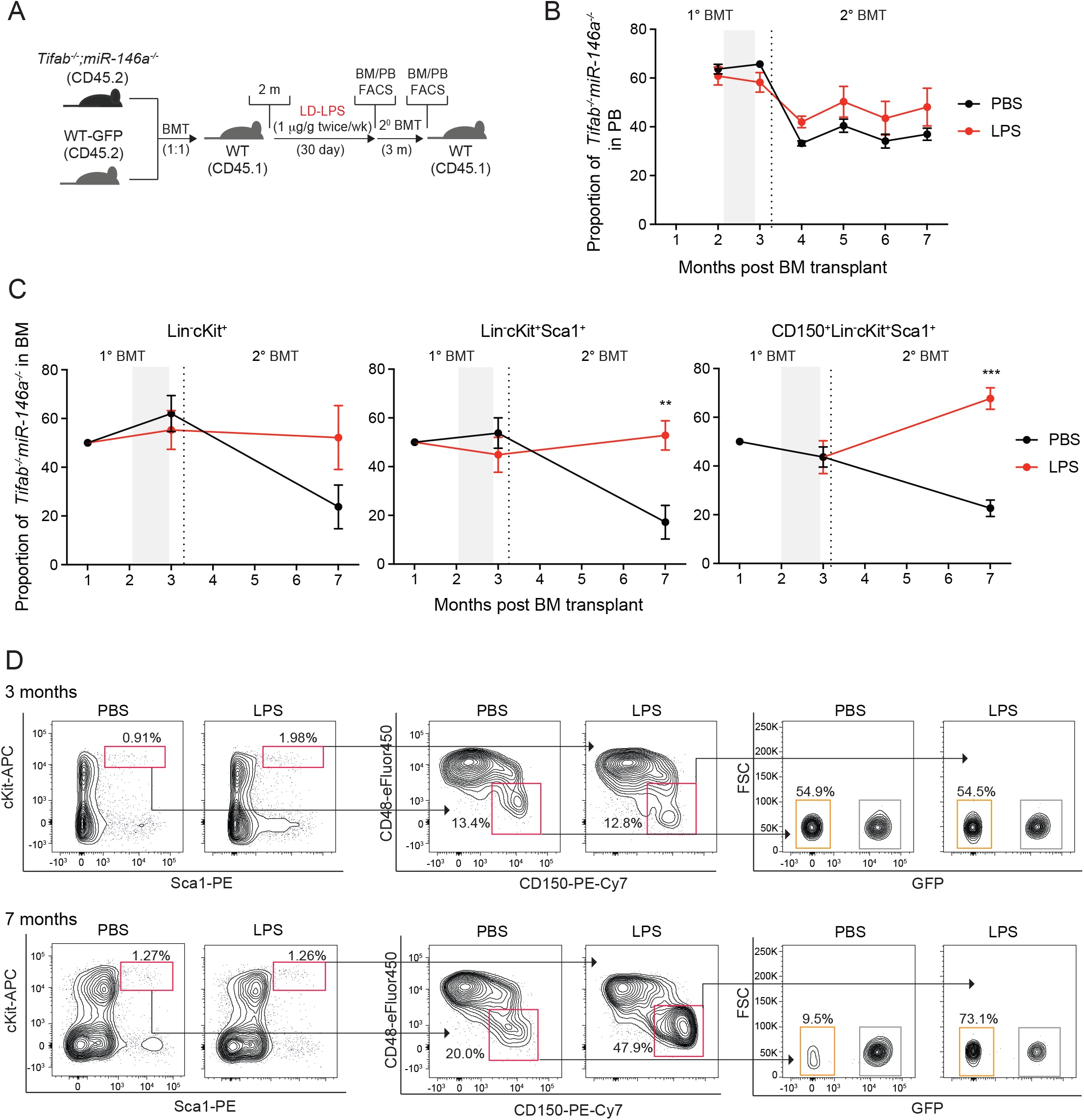
Tifab^−/-^;miR-146a^−/-^ BM HSPCs have a short-term competitive disadvantage with LD-LPS. **(A)** Outline of competitive BM transplantations using WT-GFP and Tifab^−/-^;miR-146a^−/-^ mice in the presence of low-dose LPS (LD-LPS). **(B)** Chimerism of Tifab^−/-^;miR-146a^−/-^ cells (GFP^−^ cells) in PB at the indicated time points (n = 5-6 per group). Gray bar indicates in vivo LPS treatment. Data represent the mean ± s.e.m. **(C)** Chimerism of Tifab^−/-^;miR-146a^−/-^ cells (GFP^−^ cells) in LK, LSK and HSC in the BM at the indicated time points (n = 5-6 per group). Gray bar indicates in vivo LPS treatment. Data represent the mean ± s.e.m. **(D)** Representative flow cytometric profiles of Tifab^−/-^;miR-146a^−/-^ cells (GFP^−^ cells) in HSC at 12-16 weeks after primary (upper panel) and secondary transplantation (lower panel). Significance for panels C was determined with a Student’s *t* test (**, *P* < 0.01; ***, *P* < 0.001).

### Low-grade inflammation results in impaired *Tifab*^*-/-*^*;miR-146a*^*-/-*^ HSPCs

To determine the effects of LD-LPS directly on the function of *Tifab*^*-/-*^*;miR-146a*^*-/-*^ HSPCs, we first isolated LSKs from WT and of *Tifab*^*-/-*^*;miR-146a*^*-/-*^ mice treated with in vivo chronic LD-LPS injection, and performed hematopoietic progenitor cell assays in methylcellulose. LD-LPS had minimal effects on WT LSKs as compared to untreated cells but resulted in a significant reduction of colony formation by *Tifab*^*-/-*^*;miR-146a*^*-/-*^ LSKs (**Fig 3A**). We next investigated the consequences of chronic low-grade inflammation on hematopoiesis and the long-term function of *Tifab*^*-/-*^*;miR-146a*^*-/-*^ HSPCs in vivo. Total BM cells isolated from CD45.2 *Tifab*^*-/-*^*;miR-146a*^*-/-*^ or CD45.2 WT mice injected i.p. with LD-LPS twice a week for 30 days were washed to remove residual LPS and serially transplanted every 4 months with CD45.1 WT competitor BM cells into lethally-irradiated CD45.1 WT mice (**Fig. 3B**). Exposure to LD-LPS resulted in reduced proportion of CD45.2^+^ *Tifab*^*-/-*^*;miR-146a*^*-/-*^ in the PB compared to PBS-derived CD45.2^+^ *Tifab*^*-/-*^*;miR-146a*^*-/-*^ PB cells (**Fig. 3C**). The reduction in PB chimerism is mainly due to impaired lymphopoiesis of *Tifab*^*-/-*^*;miR-146a*^*-/-*^ cells exposed to LD-LPS (**Fig. 3D**). At this time point, the production of myeloid cells in the PB was increased for *Tifab*^*-/-*^*;miR-146a*^*-/-*^ cells exposed to LD-LPS as compared to the control PBS groups following the primary BM transplantation (**Fig. 3D**), but this effect was not observed following the secondary BM transplantation (**Fig. 3E**). Exposure of CD45.2^+^ WT HSPCs to LD-LPS resulted in negligible effects at month 3 upon primary BM transplantation (**Fig. 3F**). However, exposure to LD-LPS resulted in reduced proportion of CD45.2^+^ WT cells in PB and CD45.2^+^ WT CD48^−^CD150^−^LSK (LT-HSC) in the BM compared to PBS-derived CD45.2^+^ WT cells at month 3 of the secondary BM transplantation (**Fig. 3E, G and H**), consistent with previous reports^37^. In stark contrast, CD45.2^+^ *Tifab*^*-/-*^*;miR-146a*^*-/-*^ BM cells exposed to LD-LPS resulted in significantly reduced proportions of CD45.2^+^ WT cells in the PB compared to PBS-derived CD45.2^+^ BM cells following the primary BM transplantation (**Fig 3E**). Furthermore, the *Tifab*^*-/-*^*;miR-146a*^*-/-*^ BM cells exposed to LD-LPS were unable to recover in the secondary BM transplantaton as the proportion of all HSPCs subsets were further out-competed by WT cells (**Fig. 3G-H**). These findings indicate that low-grade inflammation impairs the long-term function and self-renewal of del(5q)-like HSPCs.

**Fig. 3.**
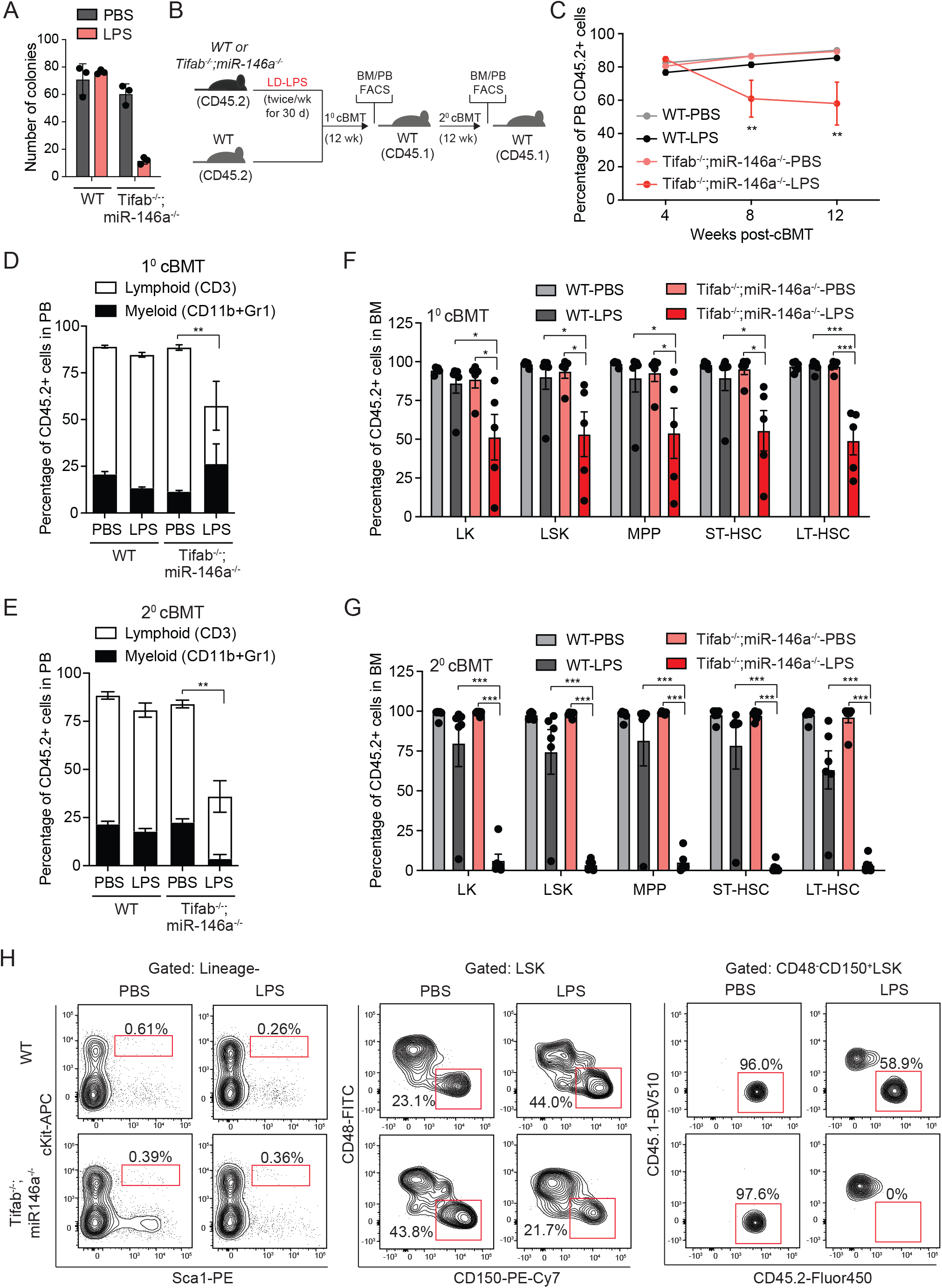
Tifab^−/-^;miR-146a^−/-^ HSPC are functionally defective following LD-LPS. **(A)** Colony assay of WT or Tifab^−/-^;miR-146a^−/-^ LSKs treated with LD-LPS (n = 3 per group). **(B)** Outline of competitive BM transplantations using WT or Tifab^−/-^;miR-146a^−/-^ mice in the presence of low-dose LPS (LD-LPS). **(C)** Summary of donor-derived PB proportions at the indicated time points (n = 5-6 per group). Error bars represent the SEM. Statistical analysis was performed between Tifab^−/-^;miR-146a^−/-^-PBS and -LPS. **(D)** Proportion of donor-derived PB cells from primary recipient mice (n = 5-6 per group). Error bars represent the SEM. **(E)** Proportions of donor-derived BM cells from primary recipient mice is reported 3 months after transplantation. Error bars represent the SEM (n = 5-6 per group). **(F)** Proportion of donor-derived PB cells from secondary recipient mice (n = 6 per group). Error bars represent the SEM. **(G)** Proportions of donor-derived BM cells from secondary recipient mice is reported 3 months after transplantation. Error bars represent the SEM (n = 6 per group). Significance for panels A, C, E, and F was determined with a Student’s *t* test (*, *P* < 0.05; **, *P* < 0.01; ***, *P* < 0.001).

One of the consequences of low-grade inflammation on HSCs is reduced quiescence due to increased cell proliferation^37,38^. Therefore, we next evaluated the effect of LD-LPS on the cell cycle status of *Tifab*^*-/-*^*;miR-146a*^*-/-*^ HSCs. *Tifab*^*-/-*^*;miR-146a*^*-/-*^ or WT mice were injected with PBS or LD-LPS (1 μg/g) i.p. twice a week for 30 days along with the administration of BrdU via drinking water and then followed by cell cycle analysis of BM cells (**Fig. 4A**). LD-LPS resulted in significantly reduced number of *Tifab*^*-/-*^*;miR-146a*^*-/-*^ HSCs in the BM as compared to PBS-treated *Tifab*^*-/-*^*;miR-146a*^*-/-*^ mice or WT mice treated with LD-LPS (**Fig. 4B**). The reduction in *Tifab*^*-/-*^*;miR-146a*^*-/-*^ HSCs in the BM treated with LD-LPS correlated with an increased frequency of BrdU+ *Tifab*^*-/-*^*;miR-146a*^*-/-*^ HSCs cells treated with LD-LPS compared to WT BM cells treated with or without LPS and untreated *Tifab*^*-/-*^*;miR-146a*^*-/-*^ BM cells (**Fig. 4B**,**C**). The reduction in *Tifab*^*-/-*^*;miR-146a*^*-/-*^ HSCs in the BM after LD-LPS was not due to decreased cell viability. The percent of apoptotic HSCs (7AAD/AnnexinV) from *Tifab*^*-/-*^*;miR-146a*^*-/-*^ mice treated with LD-LPS was similar to untreated mice or to WT treated with LD-LPS (**Fig. 4D**). Collectively, these results suggest that deletion of *Tifab* and *miR-146a* reduces the long-term repopulating potential of HSPC immediately following inflammation due to loss of cellular quiescence.

**Fig 4.**
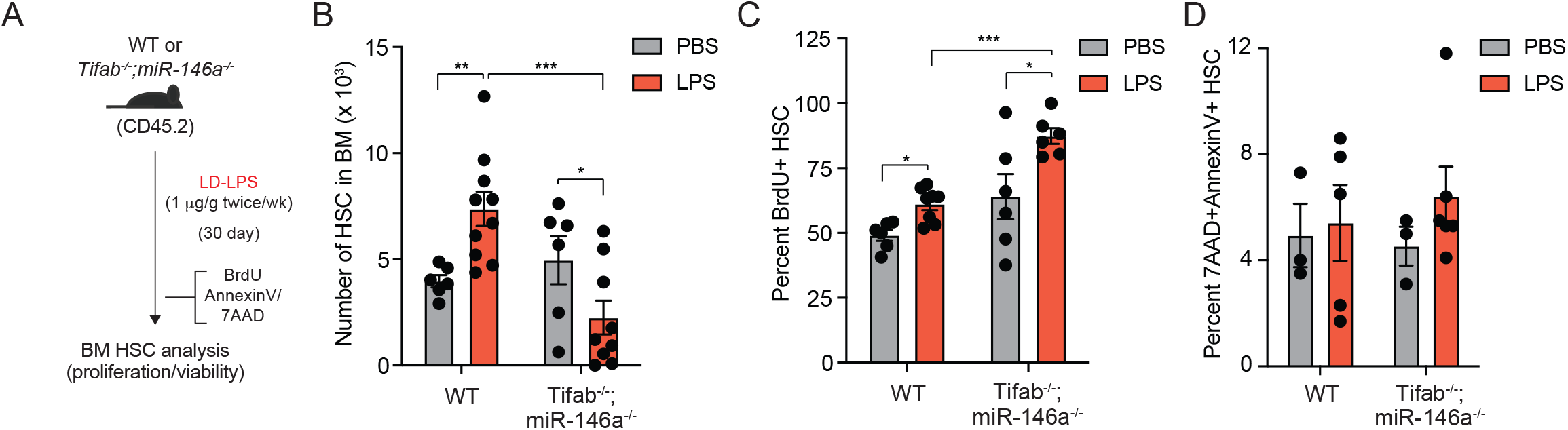
Tifab^−/-^;miR-146a^−/-^ HSPC are less quiescent following LD-LPS. **(A)** Outline of in vivo BrdU incorporation assay using WT and Tifab^−/-^;miR-146a^−/-^ mice in the presence of low-dose chronic inflammation. **(B)** Absolute number of HSC from WT and Tifab^−/-^;miR-146a^−/-^ mice treated with LPS (n=6-10). Error bars represent the SEM. **(C)** Proportion of BrdU-positive cells within HSC from WT and Tifab^−/-^;miR-146a^−/-^ mice treated with LPS (n = 6-8). Error bars represent the SEM. **(D)** Proportion of 7AAD^+^AnnexinV^+^ cells within HSC from WT and Tifab^−/-^;miR-146a^−/-^ mice treated with LPS (n = 3-5). Error bars represent the SEM. Significance for panels B, C, and D was determined with a Student’s *t* test (*, *P* < 0.05; **, *P* < 0.01; ***, *P* < 0.001).

### Low-grade inflammation results in p53 expression and activation in *Tifab*^*-/-*^*;miR-146a*^*-/-*^ HSPCs

To identify the molecular basis for the impaired function of *Tifab*^*-/-*^*;miR-146a*^*-/-*^ HSPC during low-grade inflammation, we performed RNA-sequencing in *Tifab*^*-/-*^*;miR-146a*^*-/-*^ and WT LSK treated with LD-LPS in vitro for 90 min (**Fig. 5A**). LPS treatment resulted in expression of 90 differentially upregulated genes and 212 downregulated genes in *Tifab*^*-/-*^*;miR-146a*^*-/-*^ LSK cells compared to untreated cells (**Fig. 5B**,**C** and **Supplemental Table 1**). In contrast, LPS treatment resulted in 79 differentially upregulated and 125 downregulated genes in WT LSK cells compared to untreated cells (**Fig. 5B**,**C** and **Supplemental Table 2**). Moreover, there was minimal overlap in the identify of differentially expressed genes in LPS-treated *Tifab*^*-/-*^*;miR-146a*^*-/-*^ LSK and WT LSK cells (**Fig. 5C**). Pathway analysis of differentially expressed genes using the Reactome Database revealed changes in megakaryocyte/platelet- and nicotinic acetylcholine receptor-related pathways in *Tifab*^*-/-*^*;miR-146a*^*-/-*^ LSK cells (**Fig. 5D**). WT cells treated with LPS exhibited enrichment in pathways related to epigenetic regulation, including histone methylation and acetylation (**Fig. 5D**). These results indicate TLR stimulation to *Tifab*^*-/-*^*;miR-146a*^*-/-*^ and WT HSPC results in a distinct effect on gene expression programs, pathways, and cellular states.

**Fig 5.**
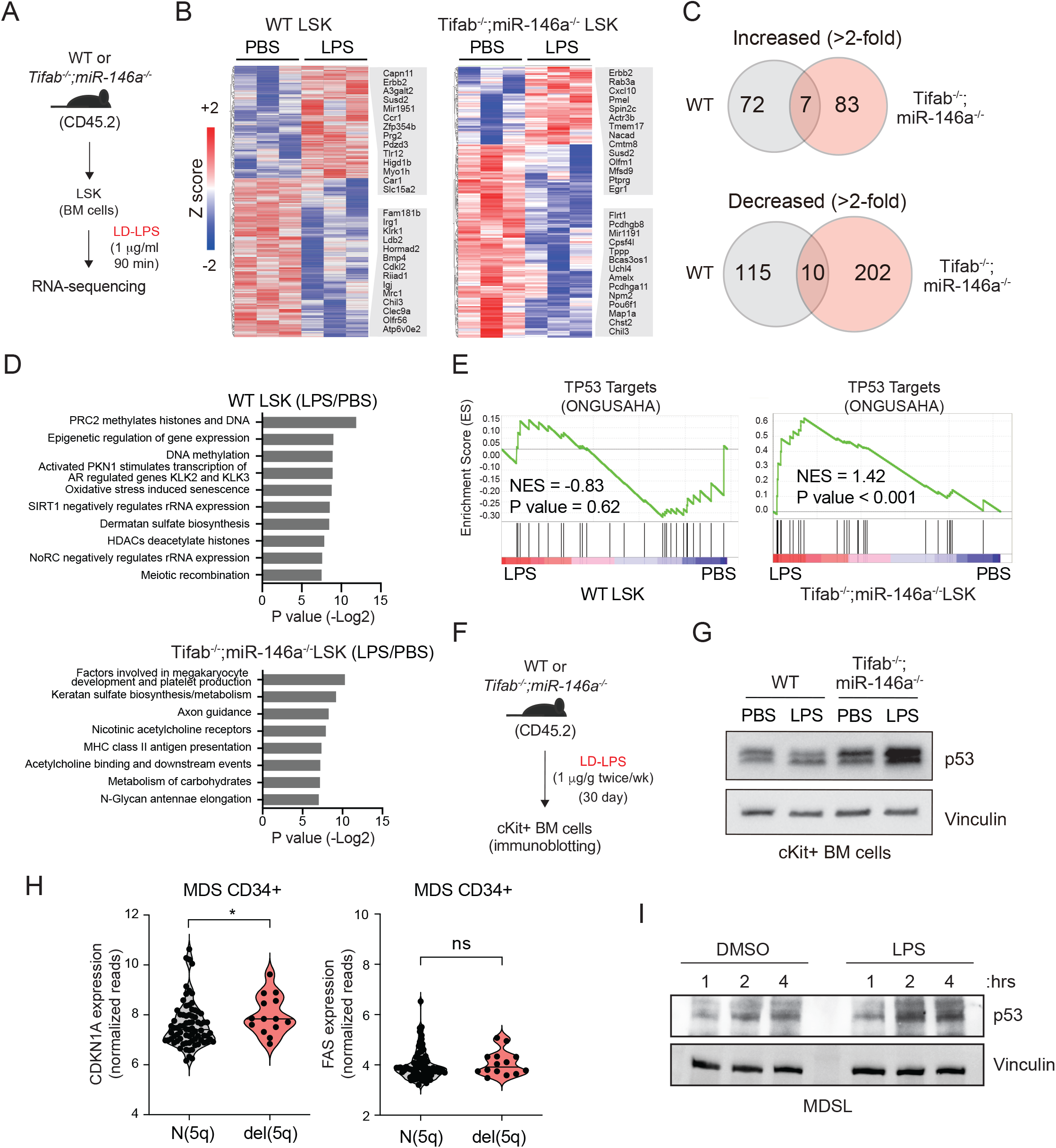
Differential TLR4 stimulation and p53 activation in *Tifab*^*-/-*^*;miR-146a*^*-/-*^ HSPCs with LD-LPS. **(A)** Outline of RNA-sequencing using HSPC from WT and Tifab^−/-^;miR-146a^−/-^ mice in the presence of low-dose LPS. **(B)** Heatmap of differentially expressed genes in WT or Tifab^−/-^;miR-146a^−/-^ LSK cells treated with LPS or PBS (1.5-fold, P < 0.05, n = 3 per group). **(C)** Venn diagram of upregulated/downregulated genes (1.5-fold, P < 0.0.5) in WT (relative to WT LSK cells treated with PBS) or Tifab^−/-^;miR-146a^−/-^ LSK cells (relative to Tifab^−/-^;miR-146a^−/-^ LSK cells treated with PBS). **(D)** Pathway analysis of LPS-stimulated WT LSK cells (relative to WT LSK cells treated with PBS) and LPS-stimulated Tifab^−/-^;miR-146a^−/-^ LSK cells (relative to Tifab^−/-^;miR-146a^−/-^ LSK cells treated with PBS). **(E)** GSEA plots for TP53 targets in LPS-stimulated WT LSK cells (relative to WT LSK cells treated with PBS) and LPS-stimulated Tifab^−/-^;miR-146a^−/-^ LSK cells (relative to Tifab^−/-^;miR-146a^−/-^ LSK cells treated with PBS). NES, normalized enrichment score. **(F)** Outline of in vivo administration of LPS following by immunoblot analysis. **(G)** Immunoblot analysis of WT and Tifab^−/-^;miR-146a^−/-^ c-Kit+ BM cells from WT and Tifab^−/-^;miR-146a^−/-^ mice treated with LD-LPS (1 μg /g) or vehicle twice a week for 30 days. **(H)** CDKN1A and FAS mRNA levels obtained from publicly-available data (GSE58831) on CD34+ cells isolated from del(5q) MDS versus normal karyotype MDS. **(I)** Immunoblot analysis of a patient-derived del(5q) MDS cell lines (MDSL) was treated with LPS (100 ng/mL) for the indicated time points.

We previously reported that TIFAB directly binds USP15, a deubiquitinating enzyme, and mediates its catalytic function by augmenting the deubiquitination of USP15 substrates. One of the key substrates regulated by the TIFAB-USP15 axis includes p53^31^. Deletion of *TIFAB* sensitizes hematopoietic cells to a variety of cellular stressors, such as 5-fluorouracil, ionizing radiation, and viral infection, which is dependent on p53 activation^31^. Moreover, p53 is implicated in inflammatory responses and is critical for maintaining homeostasis of normal HSPCs by regulating cellular self-renewal, proliferation, and survival^39,40^. Collectively, these findings prompted us to assess p53 activation in *Tifab*^*-/-*^*;miR-146a*^*-/-*^ cells exposed to low-grade inflammation. Gene set enrichment analysis (GSEA) revealed that p53-related gene signatures were positively enriched in *Tifab*^*-/-*^*;miR-146a*^*-/-*^ LSK cells treated with LPS compared to untreated *Tifab*^*-/-*^*;miR-146a*^*-/-*^ cells, while there was no significant enrichment of p53-related signature in WT cells treated with LPS (**Fig. 5E**). Baseline expression of p53-related genes were comparable between untreated WT and *Tifab*^*-/-*^*;miR-146a*^*-/-*^ HSPCs (not shown), suggesting that maximal p53 activation occurs during inflammation in del(5q)-like MDS HSPCs. To evaluate p53 protein levels in *Tifab*^*-/-*^*;miR-146a*^*-/-*^ HSPC exposed to chronic inflammation, we purified cKit+ HSPCs from WT or *Tifab*^*-/-*^*;miR-146a*^*-/-*^ mice that were treated with LD-LPS for 30 days or PBS (**Fig. 5F**). Baseline p53 protein levels were slightly increased in *Tifab*^*-/-*^*;miR-146a*^*-/-*^ HSPCs as compared to untreated *Tifab*^*-/-*^*;miR-146a*^*-/-*^ HSPCs or WT HSPCs (**Fig. 5G**). Importantly, p53 protein levels were further increased upon treatment with LD-LPS in *Tifab*^*-/-*^*;miR-146a*^*-/-*^ HSPCs as compared to untreated *Tifab*^*-/-*^*;miR-146a*^*-/-*^ HSPCs or LPS-treated WT HSPCs (**Fig. 5G**). To determine whether loss of TIFAB and miR-146a in human MDS HSPCs exhibit activation of p53, we first examined global p53-related gene signatures in MDS CD34+ with del(5q) versus ones with normal 5q status (N(5q)). p53-related signatures were not significantly different in del(5q) MDS CD34+ cells as compared to N(5q) MDS CD34+ cells. Upon closer examination of p53 target genes, expression of *CDKN1A*, a critical gene related to cell cycle regulation, was significantly increased in del(5q) MDS CD34+ cells as compared to N(5q) MDS CD34+ cells (**Fig 5H**). However, p53 target genes implicated in apoptosis (i.e., *FAS* and *BAX*) were not differentially expressed (**Fig. 5H**). This observation suggests that p53 priming in del(5q) MDS CD34+ cells is related to loss of cell cycle control rather than apoptosis induction. Moreover, LPS stimulation of a human MDS cell line resulted in a significant increase in p53 protein levels (**Fig. 5I**). Collectively, these findings suggest that del(5q) MDS HSPCs are primed to activate p53 signaling and select target genes.

### Deletion of p53 restores the functional defect of *Tifab*^*-/-*^*;miR-146a*^*-/-*^ HSPCs during low-grade inflammation

*TP53* mutations are highly enriched in del(5q) AML patients following an initial MDS diagnosis^41^; therefore, increased p53 signaling in del(5q) MDS HSPCs due to low-grade inflammation may create a selective pressure that eventually leads to genetic inactivation of p53 in the MDS HSPCs. To examine the role of p53 in *Tifab*^*-/-*^*;miR-146a*^*-/-*^ HSPCs during low-grade inflammation, we generated *Tifab*^*-/-*^*;miR-146a*^*-/-*^ mice in which one copy of *p53* is deleted (hereafter *Tifab*^*-/-*^*;miR-146a*^*-/-*^;*p53*^+/-^) (**Supplementary Fig. 3**). WT, *Tifab*^*-/-*^*;miR-146a*^*-/-*^ or *Tifab*^*-/-*^*;miR-146a*^*-/-*^;*p53*^+/-^ mice were treated with LD-LPS (1 μg/g) i.p. or PBS twice a week for 30 days (**Fig. 6A**). After 30 day exposure to LD-LPS, we confirmed that p53 expression was suppressed in *Tifab*^*-/-*^*;miR-146a*^*-/-*^;*p53*^+/-^ HSPCs as compared to *Tifab*^*-/-*^*;miR-146a*^*-/-*^ HSPCs (**Fig. 6B**). Following the last treatment with LD-LPS, BM cells (CD45.2) were isolated and co-transplanted with WT competitor BM cells (CD45.1) (one to one ratio) into lethally-irradiated CD45.1 WT mice (**Fig. 6A**). At 5 months post-transplant, LD-LPS treatment resulted in reduced proportion of *Tifab*^*-/-*^*;miR-146a*^*-/-*^ cells in the PB due to impaired production of myeloid cells (**Fig. 6C-D**), as observed in **Fig. 1C** and **Fig. 3C**,**D**. In contrast, deletion of p53 restored the PB chimerism of *Tifab*^*-/-*^*;miR-146a*^*-/-*^;*p53*^+/-^ myeloid cells exposed to LD-LPS (**Fig. 6C-D**). Furthermore, *Tifab*^*-/-*^*;miR-146a*^*-/-*^;*p53*^+/-^ HSCs did not decrease in the BM upon LD-LPS treatment as compared to untreated *Tifab*^*-/-*^*;miR-146a*^*-/-*^;*p53*^+/-^ HSCs, or LD-LPS treated *Tifab*^*-/-*^*;miR-146a*^*-/-*^ HSCs (**Fig. 6E**). Since we found that increased cell proliferation correlated with loss of *Tifab*^*-/-*^*;miR-146a*^*-/-*^ HSCs exposed to LD-LPS (**Fig. 4C**) and that del(5q) MDS CD34+ cells exhibit dysregulation of p53 target genes related to cell cycle regulation (**Fig 5H**), we performed in vivo BrdU incorporation assays. We observed that there were fewer BrdU+ *Tifab*^*-/-*^*;miR-146a*^*-/-*^;*p53*^+/-^ HSCs upon LD-LPS treatment as compared to *Tifab*^*-/-*^*;miR-146a*^*-/-*^ HSCs (**Fig. 6F**), suggesting that p53 is unexpectedly responsible for the excessive proliferation of *Tifab*^*-/-*^*;miR-146a*^*-/-*^ HSCs exposed to low-grade inflammation. To determine whether loss of p53 in *Tifab*^*-/-*^*;miR-146a*^*-/-*^ BM cells can initiate leukemia development, we aged mice for up to one year. Although deletion of p53 provides a competitive advantage of *Tifab*^*-/-*^*;miR-146a*^*-/-*^ HSPCs during low-grade inflammation, we did not observe overt AML in these mice (data not shown). This finding is consistent with observations in del(5q) MDS patients; TP53 mutations occur in low-risk del(5q) MDS at a frequency of 20% and are linked to clonal evolution and disease progression^42^. Therefore, we posit that p53 mutations preserve the competitive advantage of del(5q) HSPCs during chronic low-grade inflammation and that additional cooperating mutations contribute to the development of overt AML in del(5q) MDS.

**Fig 6.**
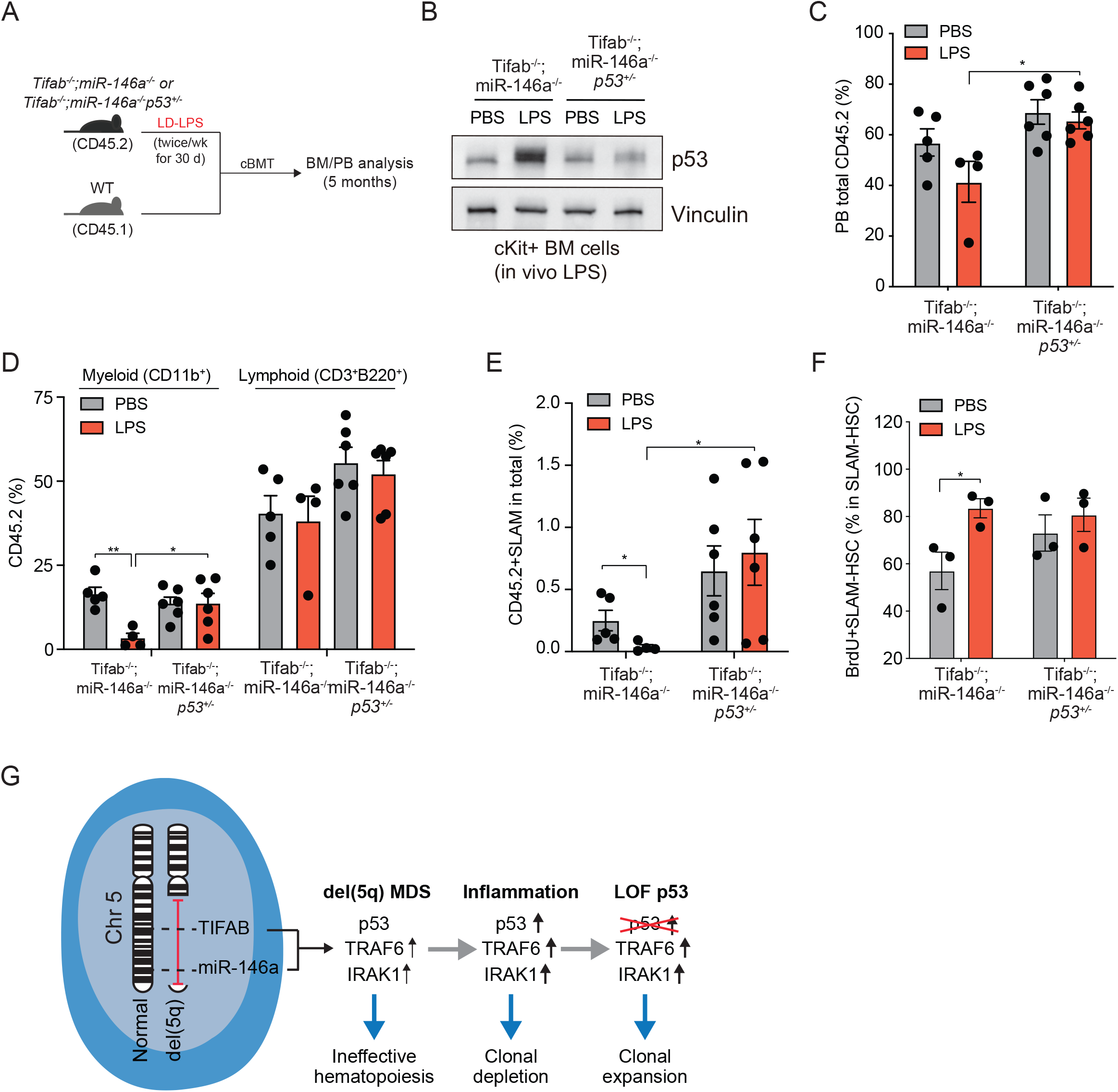
Deletion of p53 restores exhaustion of *Tifab*^*-/-*^*;miR-146a*^*-/-*^ HSCs following LD-LPS. **(A)** Outline of competitive BM transplantations using Tifab^−/-^;miR-146a^−/-^ or Tifab^−/-^;miR-146a^−/-^;p53^+/-^ mice in the presence of low-dose chronic inflammation. **(B)** Immunoblot analysis of Tifab^−/-^;miR-146a^−/-^ or Tifab^−/-^;miR-146a^−/-^;p53^+/-^ c-Kit+ BM cells from WT and Tifab^−/-^;miR-146a^−/-^ mice treated with LD-LPS (1 μg /g) or vehicle twice a week for 30 days. **(C-D)** Proportion of donor-derived PB cells from the recipient mice transplanted with Tifab^−/-^;miR-146a^−/-^ or Tifab^−/-^;miR-146a^−/-^;p53^+/-^ mice treated with either PBS or LPS. Error bars represent the SEM. **(E)** Proportion of donor-derived SLAM HSC in total cells from the recipient mice transplanted with Tifab^−/-^;miR-146a^−/-^ or Tifab^−/-^;miR-146a^−/-^;p53^+/-^ mice treated with either PBS or LPS. Error bars represent the SEM. **(F)** Proportion of BrdU-positive cells within HSC from Tifab^−/-^;miR-146a^−/-^ or Tifab^−/-^;miR-146a^−/-^;p53^+/-^ mice treated with LPS (n=3 per group). Error bars represent the SEM. **(G)** Summary of findings. Del(5q) MDS HSPCs (blue) exhibit impaired hematopoiesis and increased innate immune signaling because of derepression of TRAF6 and IRAK1 following deletion of miR-146a and TIFAB. Loss of TIFAB also results in increased p53 activation due to diminished USP15 activation. During systemic inflammation, del(5q) MDS HSCPs activated IRAK-TRAF6 signaling and induce p53, which results in clonal depletion. Loss of function mutations (LOF) or deletion of p53 permits clonal expansion of del(5q) MDS HSPCs during systemic inflammation. Significance for panels C, D, E, and F was determined with a Student’s *t* test (*, *P* < 0.05).

## Discussion

Immune dysfunction and altered innate immune signaling is a hallmark of MDS. Systemic inflammation has also been shown to favor the expansion of pre-leukemic or MDS HSCs over normal HSCs and contribute to the pathogenesis of MDS. Based on cytokine profiling of patients with pre-leukemic conditions (ICUS and CCUS) and MDS, inflammatory cytokine signatures are involved in the early stages of pathogenesis, well before the onset of MDS^43^. Several trials are currently testing therapies that interfere with the immune cell dysfunction, systemic inflammation, and altered innate immune signaling in MDS^44^. Our study investigated the effects of low-grade chronic inflammation on the function of del(5q)-like mouse MDS HSPCs and contribution to disease. Utilizing a del(5q)-like MDS mouse model in which miR-146a and TIFAB are co-deleted, we found that low-grade inflammation contributes to a fatal BM failure, a competitive advantage to functionally defective HSPCs, and p53-mediated attrition of HSPCs. These findings provide a potential explanation for the high rate of TP53 mutations in patients with del(5q).

The prevalence of TP53 mutations in all MDS subtypes is low, averaging 5–10%, but is highly associated with isolated del(5q) or complex karyotypes with -5/5q-.^41^ TP53 mutations are a predictor of poor prognosis in both de novo and secondary MDS and are associated with higher risk disease, p53 protein overexpression, increased blast count and AML progression^41^. An isolated del(5q) in MDS has a better prognosis as compared to the other MDS cytogenetic subtypes. However, *TP53* mutations occur in lower-risk del(5q) MDS at a frequency of approximately 20% and are associated with a dismal prognosis^41^. Disease progression of lower-risk del(5q) MDS is related to the evolution of pre-existing or emerging subclones carrying a *TP53* mutation^42^. *TP53* mutations at diagnosis are predictive of disease progression in lower risk del(5q) MDS and disease progression mostly occurred in patients with *TP53* mutations emerging before or at the time of progression. However, it was confirmed the absence of a TP53 mutation at diagnosis suggesting that the TP53 mutations are secondary events in del(5q) MDS patients.

The p53 tumor suppressor has important functions in cell cycle control, apoptosis, senescence, DNA repair, and genomic stability^45^. p53 can be activated by inflammation and interfere with NF-κB signaling in a variety of cell types^46^. It is also well established that p53 mutations are acquired in hematopoietic cells in patients receiving genotoxic therapy, leading to therapy-related MDS or AML^47^. In this scenario, the genotoxic stress is a selective pressure for loss of p53-mediated apoptosis in the hematopoietic cells. However, the reasons for the high frequency of p53 mutations in del(5q) and which of the p53 functions are essential in del(5q) MDS is insufficiently understood. Moreover, it also remains unresolved why these p53 functions are selected against during clonal evolution and disease progression. In a related BMF syndrome, Fanconi anemia, chronic inflammation and p53 activation contribute to the clinical manifestation of disease^48^. In contrast to our data herein, p53 activation in FA results in profound cell cycle arrest. Based on our in vitro and in vivo models, we propose that loss of p53-mediated cell cycle control of MDS HSPCs is the primary p53 function selected against during inflammation (**Fig. 6G**). Paradoxically, we find that elevated p53 levels in del(5q)-like MDS HSCPs following inflammation correlate with increased HSC proliferation and a reduced competitive advantage, while loss of p53 in these cells results in reduced HSC proliferation and an increased competitive advantage. This paradoxical observation may be explained by the unique dependency of p53 in the context of del(5q)-like MDS HSPCs.

Deletion of miR-146a and TIFAB as in del(5q) MDS contribute to altered responses to systemic inflammation and among the many changes, in increased p53 activation in HSPCs. Consequently, systemic inflammation results in fewer number and more proliferative haploinsufficient miR-146a and TIFAB HSCs. To overcome the negative effects of systemic inflammation, we posit that haploinsufficient miR-146a and TIFAB HSCs have a proclivity to gain a competitive advantage upon loss of p53, as our data suggests (see Fig. 6). TIFAB has previously been implicated in p53 regulation by directly regulating USP15 ubiquitin hydrolase activity^31^. Expression of TIFAB in HSPCs permits USP15 signaling to substrates, including MDM2, and mitigates p53 expression in leukemic myeloid cells. As such, TIFAB-deficient HSPCs exhibit compromised USP15 signaling and are sensitized to a variety of hematopoietic stressors by derepression of p53. Thus, our finding that loss of miR-146a and TIFAB in HSCs results in increased p53 expression during inflammation is likely due to a cooperative effect through both del(5q) genes. Interestingly, deletion of p53 in haploinsufficient miR-146a and TIFAB HSCs did not result in overt leukemia, suggesting that the primary consequence of p53 loss is to provide del(5q) MDS HSCs a competitive advantage during system inflammation. Alternatively, additional genes on the deleted region of chr 5q cooperate with loss of p53 to manifest overt leukemia. Indeed, deletion of EGR1, a del(5q) gene and regulator of p53-dependent functions, leads to myeloid leukemias upon treatment of mice with DNA alkylating agents^49^ As the current study focused on two del(5q)-related genes, miR-146a and TIFAB, it likely that additional genes within the deleted segment on chr 5q also contribute to the effects of systemic inflammation in del(5q) MDS. Future studies that take into consideration additional del(5q)-related genes are warranted.

Increased innate immune signaling in del(5q) MDS HSPCs is attributed to several haploinsufficient genes, including miR-143, miR-145, miR-146a, mDia1, and TIFAB. The individual contribution of each of these haploinsufficent genes to MDS phenotypes has been extensively evaluated in mouse models^16,17^. More recently, various models were generated to examine the cooperation of multiple haploinsufficent del(5q) genes. Mice lacking miR-143 and miR-145 have impaired HSPC activity with depletion of functional HSCs, but activation of progenitor cells^50^. Lam et al identified components of the transforming growth factor β (TGFβ) pathway as key targets of miR-143 and miR-145. As expected, the combined deletion of miR-146a with RPS14 and CSNK1A1 recapitulated many cardinal features of the 5q-syndrome, including more severe anemia with faster kinetics than Rps14 haploinsufficiency alone and pathognomonic megakaryocyte morphology^14^. Combined hematopoietic-specific deletion of Tifab and miR-146a resulted in more rapid and severe cytopenia, and progression to a fatal BM failure-like disease as compared with Tifab- or miR-146a-deficiency alone^24^. HSPCs from Tifab^−/-^, miR-146a^−/-^ and dKO mice exhibit enrichment of gene regulatory networks associated with innate immune signaling. Moreover, a subset of the differentially expressed genes is controlled synergistically following deletion of Tifab and miR-146a^24^. Nearly half of these defined synergy response genes identified in the mouse models were aberrantly expressed in del(5q) MDS HSPC when TIFAB and miR-146a were both deleted. Dual deficiency of mDia1 and miR-146a caused an age-related anemia and ineffective erythropoiesis mimicking human MDS^51^. It was also demonstrated that the aging BM microenvironment was important for the development of ineffective erythropoiesis in these mice. Damage-associated molecular pattern molecules (DAMPs), whose levels increase in aging BM, induced TNFα and IL-6 upregulation in myeloid-derived suppressor cells (MDSCs) in the mDia1/miR-146a double knockout mice^51^. Of note, loss of miR-146a alone is sufficient to induce HSC defects and hematopoietic disease^52,53^. Deletion of miR-146a in HSCs promotes premature HSC aging and inflammation, preceding development of aging-associated myeloid malignancy. These effects are mediated in part by excessive signaling through its targets TRAF6 and IRAK1^21,54,55^. The mechanistic basis for the altered response and functional impairment of miR-146a/TIFAB-deficient HSCPs during inflammation requires additional exploration. Interestingly in separate studies, overexpression of TRAF6 provided a competitive advantage of HSPCs during systemic inflammation by altering the NF-κB response^7^. In contrast, deletion of TRAF6 in pre-leukemic HSPCs resulted in excessive HSPC proliferation, myelopoiesis, and increased MYC signaling^56^. Collectively, these findings confirm that aberrant innate immune signaling in del(5q) MDS HSCs and altered responses to the inflammatory microenvironment play a critical role in the pathogenesis and complex phenotype of myeloid malignancies.

Several attempts are being pursued to restore normal innate immune signaling in MDS HSCs^23,57-61^. One approach has been to inhibit IRAK4 in MDS^21^. There are ongoing clinical studies evaluating IRAK4 inhibitors for low-risk MDS patients, which will provide insight into the benefit of suppressing cell-intrinsic innate immune signaling pathways downstream of IRAK-TRAF6 to improve cytopenias. In our study, we utilized a novel IRAK1/4 inhibitor, which improved cyotpenias in the del(5q)-like MDS model. These findings suggest that suppressing IRAK-TRAF6 signaling can restore blood counts upon loss of miR-146a and/or TIFAB in MDS patients. Moreover, it is also possible that suppressing chronic inflammation in low-risk del(5q) MDS patients may diminish the selective pressures leading to acquired p53 mutations. Such an approach has been demonstrated experimentally wherein the selective pressure against p53 in models of lymphoma can be mitigated by targeting the apoptotic pathway^62^. Therefore, future studies should determine whether mitigating systemic inflammation in del(5q) MDS may reduce the likelihood of secondary p53 mutations and progression towards more aggressive disease. In summary, we report that low-grade inflammation accelerates the del(5q) MDS-like phenotype and provide a potential explanation for the high rate of TP53 mutations in patients with del(5q).

## Material and Methods

### Materials

LPS (L6529) and LPS-EB Ultrapure (TLRL-PEKLPS) were purchased from Sigma and InvivoGen, respectively. The IRAK1/4 inhibitor (NCGC-1481) has been previously described^33,34^. MDS-L cells have been previously described^63,64^.

### Mice

Generation of *Tifab*^−/-^;*miR-146a*^−/-^ mice was described in our previous study^24,25^. UBI-GFP mice (Jackson Laboratory, 004353) were purchased from Jackson Laboratory. *Tifab*^−/-^;*miR-146a*^−/-^ mice were crossed with *Trp53*^−/-^ mice (Jackson Laboratories, 002101). All mice were bred, housed and handled in the Association for Assessment and Accreditation of Laboratory Animal Care-accredited animal facility of Cincinnati Children’s Hospital Medical Center.

### Hematological analysis

Blood counts were measured with a hemacytometer (HEMAVET).

### BM transplantation

For non-competitive BM transplantations, 1 × 10^6^ CD45.2^+^ BM cells were transplanted into lethally-irradiated recipient mice (CD45.1^+^ B6.SJL^Ptprca Pep3b/Boy^; 6-10 weeks of age). For competitive transplantation, BM cells from 8-week-old recipient mice were transplanted into lethally irradiated recipient mice with CD45.1^+^ B6.SJL^Ptprca Pep3b/BoyJ^ BM cells or CD45.2^+^ C57BL/6-Tg(UBC-GFP)30Scha/J BM cells. For serial transplantation, BM cells were collected from all recipient mice 3 months after transplantations, pooled together, and then 2 × 10^6^ BM cells were transplanted into lethally-irradiated recipient mice (CD45.1+ B6.SJL^Ptprca Pep3b/Boy^; 6-10 weeks of age).

### Colony forming assay

Either chronic LD-LPS (1 μg/g) or PBS to WT and *Tifab*^−/-^;*miR-146a*^−/-^ mice was administered via intraperitoneal injection twice a week for 30 days, and BM cells were harvested afterwards. Twenty thousand BM cells per replicate were plated in methylcellulose (3434; Stemcell Technologies). Colonies propagated in culture were scored at day 14.

### Flow cytometry

For immunophenotypic analysis of lineage positive cells, PB samples were processed with 1 x RBC lysis buffer, and then incubated with CD11b-PE-cy7 (25-0112-81, eBiosciences), Gr1-eFluor450 (48-5931-82, eBiosciences), CD3-PE (12-0031-83, eBiosciences), and B220-APC (17-0452-82, eBiosciences). To distinguish donor from recipient hematopoietic cells, PB were stained with CD45.1-Brilliant Violet 510 (110741, BioLegend), and CD45.2-APC-eFluor780 (47-0454-82, eBiosciences) or CD45.2-eFluor450 (48-0454-82, eBiosciences). For HSC analysis, BM cells were washed and incubated for 30 minutes with biotin conjugated lineage markers (CD11b, Gr1, Ter119, CD3, B220, mouse hematopoietic lineage biotin panel, [88-7774-75 eBiosciences]), followed by staining with streptavidin eFluor780 (47-4317-82, eBiosciences), Sca-1-PE (12-5981-82, eBiosciences), c-Kit-APC (17-1171-81, eBiosciences), CD48-FITC (11-0481-85, Affymetrix), CD150-PE-cy7 (115914, BioLegend). SLAM-HSC were identified based on expression of Lin^−^ Sca^−^1^+^c-Kit^+^CD150^+^CD48^−^.

### Cell cycle and apoptosis analysis

BrdU (Sigma-Aldrich) was administered continuously to mice via drinking water (0.5 mg/ml). After 1 week, BrdU incorporation was analyzed using a BrdU Flow Kit (559619, BD Biosciences) according to the manufacture’s recommendation. Annexin V viability staining was carried out according to manufacturer’s instructions (C0556419, BD Biosciences).

### Immunoblotting

Cell extract was prepared by lysing cells in sodium dodecyl sulfate (SDS) sample buffer followed by incubation with benzonase (70746, Millipore) on ice for 10 minutes. Nuclear and cytoplasmic fractionation was performed with Nuclear Extract Kit (40010, Active Motif) according to the manufacture’s protocol. Samples were boiled at 95 °C for 5 minutes and loaded to SDS-polyacrylamide gel electrophoresis (PAGE) and transferred to nitrocellulose membranes (162-0112, Bio-Rad). Immunoblot analysis was performed with the following antibodies: p53 (2524S, Cell Signaling), Vinculin (13901, Cell Signaling).

### RNA sequencing

Total RNA was extracted from the cells using RNeasy Plus Micro Kit (Qiagen). The initial amplification step for all samples was done with the NuGEN Ovation RNA-Seq System v2. The assay was used to amplify RNA samples to create double stranded cDNA. The concentrations were measured using the Qubit dsDNA BR assay. Libraries were then created for all samples using the Illumina protocol (Nextera XT DNA Sample Preparation Kit). The concentrations were measured using the Qubit dsDNA HS assay. The size of the libraries for each sample was measured using the Agilent HS DNA chip. The concentration of the pool was optimized to acquire at least 15-20 million reads per sample. The analysis of RNA sequencing was performed with iGeak^65^. Gene set enrichment analysis (GSEA) was performed as previously described^66^.

### Statistical analysis

The number of animals, cells, and experimental/biological replicates can be found in the figure legends. Differences among multiple groups were assessed by one-way analysis of variance (ANOVA) followed by Tukey’s multiple comparison posttest for all possible combinations. Comparison of two group was performed using the Mann-Whitney test or the Student’s *t* test (unpaired, two tailed) when sample size allowed. Unless otherwise specified, results are depicted as the mean ± standard deviation or standard error of the mean. A normal distribution of data was assessed for data sets >30. For correlation analysis, Pearson correlation coefficient (r) was calculated. D’Agostino and Pearson and Shapiro-Wilk tests were performed to assess data distributions. For Kaplan-Meier analysis, Mantel-Cox test was used. All graphs and analysis were generated using GraphPad Prism software or using the package ggplot2 from R^67^.

### Data availability

The data that support the findings of this study are available from the corresponding author upon request.

## Acknowledgements

This work was supported in parts by the National Institute of Health (U54DK126108, R35HL135787, R01DK113639 to DTS), Cincinnati Children’s Hospital Research Foundation (DTS), The Uehara Memorial Foundation (TM), The Waksman Foundation of Japan (TM), The Mochida Memorial Foundation for Medical and Pharmaceutical Research (TM), Japan Society for the Promotion of Science (TM), and Ohio State University Comprehensive Cancer Center (TM). TM is a Leukemia and Lymphoma Society Special Fellow.

