## Supplemental Figures for "Chronic inflammation suppresses del(5q)-like MDS HSCs via p53"

### Supplemental Figure 1

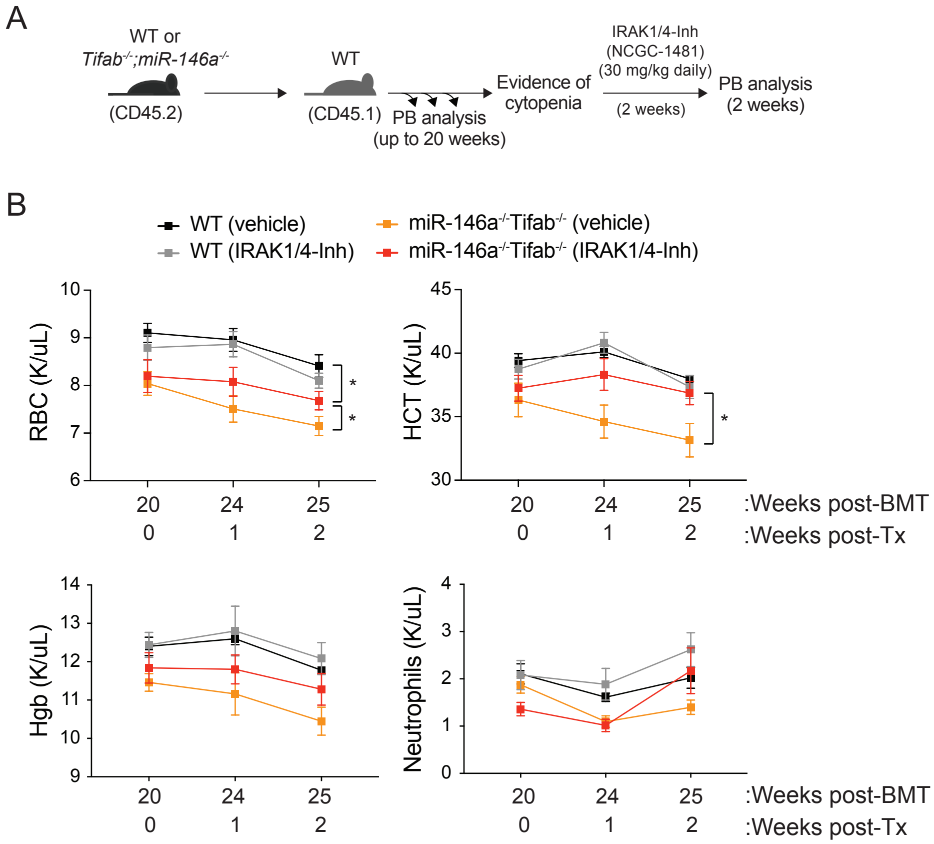


**Supplemental Figure 1. IRAK1/4-inhibitor restores blood counts in del(5q)-like MDS model. (A)** Outline of BM transplantations using WT or Tifab-/-;miR-146a-/- BM cells. PB analysis was performed monthly on recipient mice to monitor for cytopenias. At onset of cytopenias in the Tifab-/-;miR-146a-/- recipient mice, an IRAK1/4 inhibitor (NCGC-1481 at 30 mg/kg) was administered daily (or PBS, vehicle control). PB counts were performed weekly after the treatment was initiated. **(B)** PB counts of the recipient mice before (20 weeks post BM transplantation) and after treatment (post-Tx) (n = 4-5 per group). Significance for panel B was determined with a Student's t test (*, $P < 0.05$) between treated and untreated groups.

### Supplemental Figure 2.

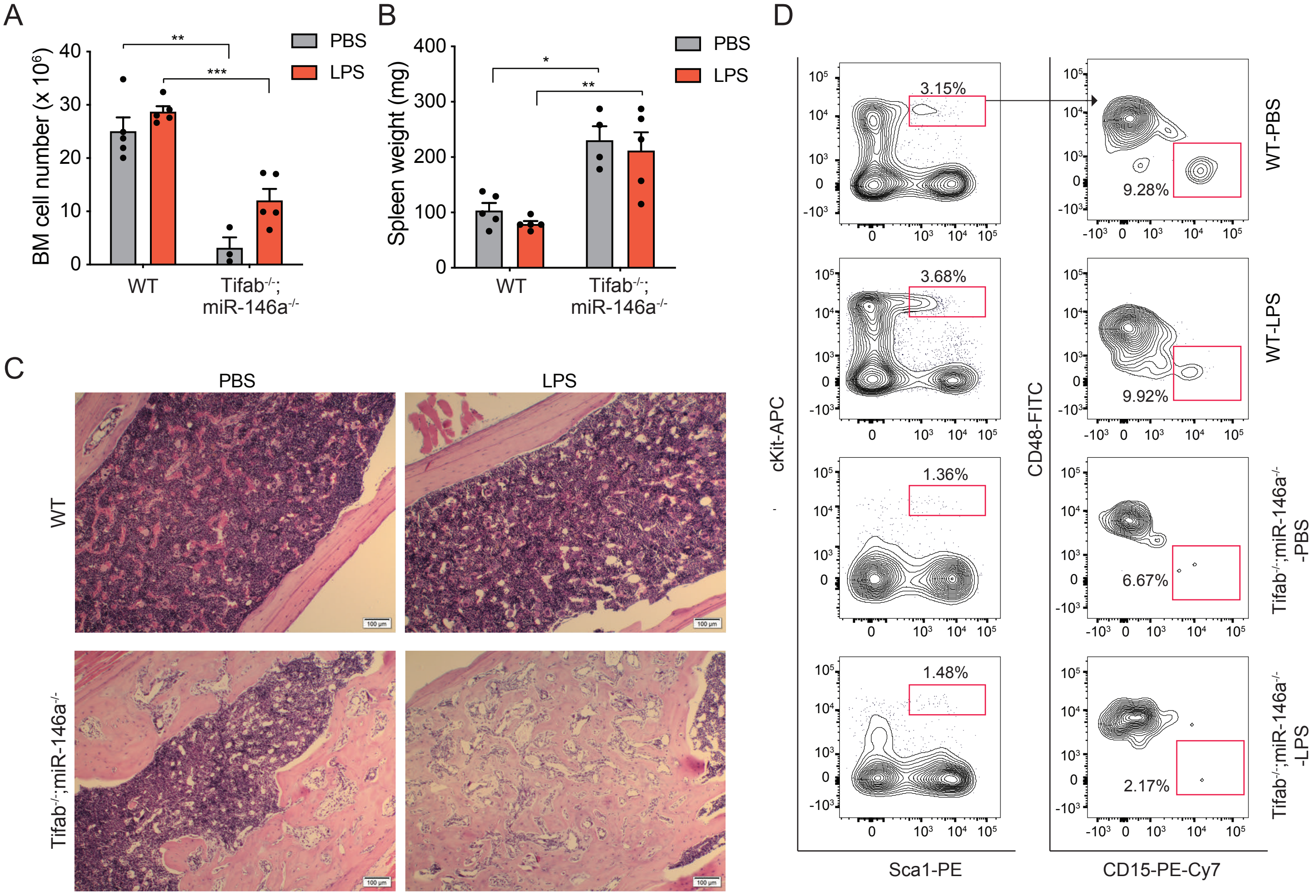


**Supplemental Figure 2. Characterization of moribund del(5q)-like MDS model. (A)** Total BM cells numbers in moribund mice transplanted with Tifab-/-;miR-146a-/- BM cells (n = 4-5 per group). WT recipient mice are age-matched controls. **(B)** Spleen weights of moribund mice. WT recipient mice are age-matched controls. **(C)** H&E staining of BM from moribund mice transplanted with Tifab-/-;miR-146a-/- BM cells (n = 4-5 per group). WT recipient mice are age-matched controls. **(D)** Representative flow cytometric analysis and gating strategy of moribund mice transplanted with Tifab-/-;miR-146a-/- BM cells. WT recipient mice are age-matched controls.

**Supplemental Figure 3.**

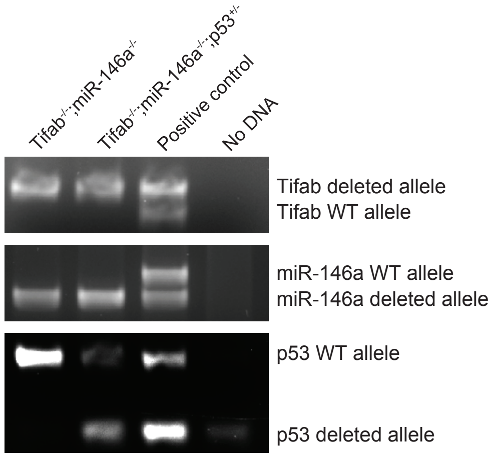


**Supplemental Figure 3. Generation of p53-deficient Tifab-/-;miR-146a-/- mice.** Genotyping analysis of p53-deficient Tifab-/-;miR-146a-/- mice.
